# The role of genetic diversity in the evolution and maintenance of environmentally-cued, male alternative reproductive tactics

**DOI:** 10.1101/528703

**Authors:** KA Stewart, R Draaijer, MR Kolasa, IM Smallegange

**Affiliations:** Department of Evolutionary Population Biology, Institute for Biodiversity and Ecosystem Dynamics, University of Amsterdam, PO Box 94240, 1090 GE, Amsterdam, The Netherlands; Institute of Systematics and Evolution of Animals, Polish Academy of Sciences, Slawkowska 17 St., 31-016 Krakow, Poland

**Author notes:** Authors declare equal first-author contribution (KA Stewart, R Draaijer). Corresponding Author: Kathryn A. Stewart.

**Keywords:** inbreeding depression, epistasis, genetic correlation, environmental threshold model, phenotypic plasticity, conditional strategy

## Abstract

**Background:** Alternative reproductive tactics (ARTs) are taxonomically pervasive strategies adopted by individuals to maximize reproductive success within populations. Even for conditionally-dependent traits, consensus postulates most ARTs involve both genetic and environmental interactions (GEIs), but to date, quantifying genetic variation underlying the threshold disposing an individual to switch phenotypes in response to an environmental cue has been a difficult undertaking. Our study aims to investigate the origins and maintenance of ARTs within environmentally disparate populations of the microscopic bulb mite, *Rhizoglyphus robini*, that express ‘fighter’ and ‘scrambler’ male morphs mediated by a complex combination of environmental and genetic factors.

**Results:** Using never-before-published individual genetic profiling, we found all individuals across populations are highly inbred with the exception of scrambler males in stressed environments. In fact within the poor environment, scrambler males and females showed no significant difference in genetic differentiation (Fst) compared to all other comparisons, and although fighters were highly divergent from the rest of the population in both poor or rich environments (e.g., Fst, STRUCTURE), fighters demonstrated approximately three times less genetic divergence from the population in poor environments. AMOVA analyses further corroborated significant genetic differentiation across subpopulations, between morphs and sexes, and among subpopulations within each environment.

*Conclusion:* Our study provides new insights into the origin of ARTs in the bulb mite, highlighting the importance of GEIs: genetic correlations, epistatic interactions, and sex-specific inbreeding depression across environmental stressors. Asymmetric reproductive output, coupled with the purging of highly inbred individuals during environmental oscillations, also facilitates genetic variation within populations, despite evidence for strong directional selection. This cryptic genetic variation also conceivably facilitates stable population persistence even in the face of spatially or temporally unstable environmental challenges. Ultimately, understanding the genetic context that maintains thresholds, even for conditionally-dependent ARTs, will enhance our understanding of within population variation and our ability to predict responses to selection.

## Background

In numerous species, it is common for individuals (usually males) to adopt different strategies to increase their reproductive success when intrasexual competition is intense. These strategies can ultimately lead to diversity within populations, comprising of characteristics such as behaviour, physiology, or morphology [1]. Referred to as alternative reproductive tactics (ARTs), strategies such as these encompass trade-offs between increased reproductive potential versus the costs incurred to produce traits under selection, often leading to the development of a less energetically demanding tactic, such as sneakers (versus guards) or satellites (versus callers). Although taxonomically widespread and studied in various organisms [1], the proximate mechanisms responsible for ART trait evolution, or the processes that maintain ARTs within single populations are not always well understood. Some ARTs are plastic by nature, driven by seemingly pure environmental effects (e.g., dung beetles, *Onthophagus acuminatus*; [2]), whereas others are fixed, determined exclusively by genetic underpinnings (e.g. lekking sandpiper, *Philomachus pugnax*; [3]), although the latter remains a relatively rarer phenomenon [4–6]. More commonly however, species demonstrating ARTs involve a combination of both genetic and environmental influences, that interrelate in genotype-by-environment interactions [7].

Genotype-by-environment interactions (GEIs) are routinely observed in traits linked to fitness [8] such that in different environments, numerous genotypes may display and switch superiority (ecological cross-over), assisting in the maintenance of variation within populations. Moreover, male sexually selected traits often show condition-dependence that is assumed to involve many loci, providing ample opportunity for mutations (‘genic capture hypothesis’ [9]) and genetic variation. For example, high genetic diversity (heterozygosity) has been linked to an individual’s fitness and condition, including an increase in survival [10, 11], parasite resistance [12], developmental stability [13], competitive ability [14], viability [9, 15, 16], mating opportunities [17], and the expression of costly secondary sexually selected traits [18]. Together, GEIs and condition-linked genetic diversity may help to reconcile the origin and maintenance of ARTs within populations [19], despite presumably strong selective forces promoting the canalization of traits, and genetic erosion associated with sexual selection (‘the lek paradox’) [19, 20].

Currently, the environmental threshold model, which links condition-dependence and GEIs [21–23], is the most widely accepted process for ART expression. Specifically, this model posits that environmental circumstances experienced by an individual during ontogeny leads to an all-or-none response in terms of expressing ARTs, which in-turn is likely influenced by the organism’s genetic background [23]. Male polymorphic variation is thus thought of as a threshold trait based on a continuously distributed phenotype that is environmentally sensitive [24]. Threshold traits have been shown to have a heritable basis, although more likely due to the heritability of the underlying threshold itself (liability traits) [25, 26]. If this polymorphic variation is under polygenic control, condition-linked genetic diversity likely plays an important role in trait expression. ARTs involve complex traits that can be heritable, subject to selection, and evolve, yet to date, the genetic basis underlying the evolution of conditionally-dependent ARTs has been difficult to quantify [27].

The bulb mite *(Rhizoglyphus robini)* is a microscopic agricultural pest, which thrives on invading crops and disperses easily when food is deprived (a species familiar with fluctuating environment conditions) [28]. This species demonstrates a short generation time, has high reproductive potential [29], and is easily reared in laboratory conditions, making it an ideal organism for experimental evolution studies. Intriguingly, the bulb mite demonstrates a complex ART system that has recently described up to three male polymorphisms, including a ‘megascrambler’ [30] that, due to its rareness within populations, will be excluded from the current study. Of the two prominent male ARTs in *R. robini*, individuals express either a ‘fighter’ or ‘scrambler’ mating tactic consisting of the ontogenetic development (or not) of weaponry comprised of a thickened, sharply terminated third pair of legs used to combat and kill rival males (fighters and scramblers, respectively) (Fig. 1). The environmental threshold model is a good candidate model to explain the evolution of this male dimorphism as high nutritional quality and quantity during development increases juvenile body size, which in turn increases fighter morph expression in adulthood [31, 32]. An experimental test of this model’s predictions on evolutionary shifts in ART expression indeed confirmed threshold shifts when selecting against fighter expression. This analysis, however, failed to capture the observed evolutionary threshold shifts when selecting against scrambler expression [33], likely because scrambler expression shifted evolutionarily in response to the demographic consequences of the experimental treatment, rather than the treatment itself. It therefore seems likely that multiple environmental drivers are involved to maintain this male dimorphism [34]. Previous research also demonstrates the bulb mite ART is somewhat heritable, yet these heritability scores vary widely depending on population or study, ranging between 0.18 to >1.00 based on experimental and modelling estimates [35–37], further suggesting this ART likely does not represents a simplistic environmental or genetic trigger.

**Figure 1.**
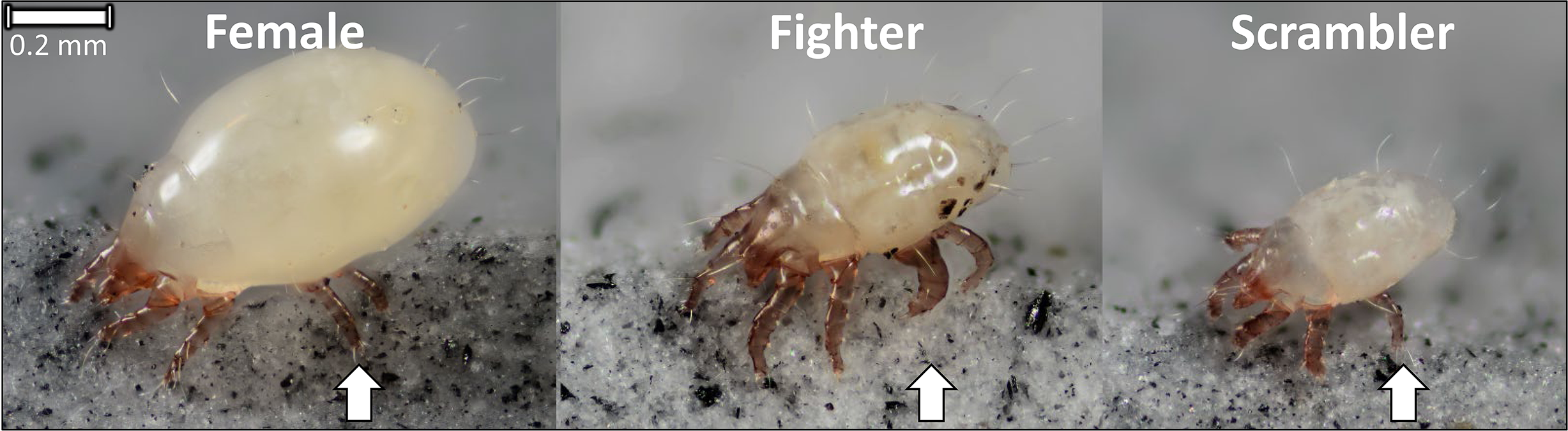
Dorsolateral photographic images of adult bulb mites (*Rhizoglyphus robini*) including the female, and male ARTs (fighter, and scrambler). All individuals are presented at the same scale (scale bar: top left) and aligned from largest to smallest (left to right), with arrows indicating major structural differences in the third-leg pair among sexes and morphs. Photographs produced by Jan van Arkel, 2017.

Here, we aim to resolve broad evolutionary questions surrounding the origin and maintenance of ARTs by quantifying the cryptic variance underpinning threshold responses to environmental cues. We do this by testing the hypothesis that genetic diversity differs between the two male ARTs in the bulb mite such that larger fighters with associative high body condition will demonstrate higher levels of genetic diversity compared to their smaller, poorer conditioned scrambler counterparts. Using populations consisting of tens-of-thousands of individual bulb mites reared under different environments, we quantified underlying genetic context in relation to ART expression, using never before published genetic markers to quantify individual-level genetic diversity across populations.

## Materials & Methods

### Specimen maintenance and collection

We used bulb mites from stock populations originating from 10 sampling sites via collecting flower bulbs near Anna Paulowna, North Holland, Netherlands in 2010, that ultimately comprise tens-of-thousands of individuals. Mites were reared and maintained at the Institute for Biodiversity and Ecosystem Dynamics at the University of Amsterdam, Netherlands in a controlled environmental chamber (25 ± 1 °C, 60% relative humidity, 16:8h light-dark photoperiod; sensu [38]) under two different rearing environments commonly used in life history studies to assess growth, development, and ART expression of mites from the family Acaridae (e.g., [39–41]). These two environments, henceforth be described as ‘poor’ or ‘rich’, differed only in their nutritional resources; mites were fed either rolled oats (poor food quality) or dried yeast (rich food quality via high quantity of protein), ad libitum. The rich resource treatment (yeast), in fact, creates a similar rearing environment to that of natural bulb mite populations feeding on garlic bulbs [39].

From the rich and poor environments, mites were randomly collected and examined with a stereomicroscope for identification. Sexes and ART morphs were identified according to the morphological criteria described by Smallegange [38], including size delimitation, genitalia, and the presence/absence of enlarged third leg pairs (main ART trait differentiation). Following recommendations that 20-30 individuals assayed within populations yield sufficiently reliable estimates for population genetic parameters [42], a total of 231 mites were sampled from the stock populations in both rich (n=126) and poor (n=105) environments, including 72 scrambler males (rich n=42, poor n=30), 76 fighter males (rich n=32, poor n=44), and 83 females (rich n=52, poor n=31). Upon collection, mites were individually preserved in 1.5 mL Eppendorf tubes containing 95% ethanol, and stored at − 20°C until DNA extraction.

Because we aimed to create equal sampling of each subpopulation (female, fighter, scrambler), the representative sex-ratio from the overall environments did not reflect the female biased operational sex-ratio from either stock or natural populations [33]. However, this sampling scheme, should bear no influence on our interpretation of whether genetic context influences the expression of ARTs in the bulb mite system.

### DNA extraction, PCR amplification, and nSSR analysis

Prior to extraction, all ethanol within the Eppendorf tubes was evaporated. For the female, and fighter male mites, we used a modified protocol from Knegt *et al.* [43]in which chelex-based DNA extraction was performed: 4-5 zirconium beads, 50μL of a 5% chelex solution (Bio-Rad laboratories), and 5μL of proteinase K (20 mg ml^-1^) were added to each tube, after which the samples were homogenized 3 times for 30s at 6500 rpm using a Precellys24 tissue homogenizer (Bertin Corp). Upon homogenization, the samples were incubated for 2 hours at 56°C, and proteinase K was inactivated via incubation for 8 minutes at 95°C. Samples were centrifuged for 2 minutes at 14000 rpm and thereafter stored at −20°C.

As scrambler male mites are typically much smaller than their fighter or female counterparts, we adjusted the DNA extraction protocol as follows: after the evaporation of all ethanol, 4.5μL proteinase K (20 mg ml^-1^) was added to each Eppendorf tube, and with a pestle, mites were ground into small pieces, after which 30μL of a 5% chelex solution was added to each tube. The samples were subsequently incubated for 3 hours at 56°C, and proteinase K was inactivated via incubation for 8 minutes at 95°C. Samples were vortexed and centrifuged shortly, and stored at −20°C prior to DNA amplification.

In total we tested 16 nuclear simple sequence repeats (nSSR) primer pairs designed for our species at Jagiellonian University in Kraków [44], optimizing the primer pairs and concomitant PCR protocol for our own populations using Dreamtaq polymerase (Thermo Fisher Scientific). Each primer pair was amplified individually in 15μL reactions wherein each reaction contained 3μL 5 X Dreamtaq buffer, 3μL dNTPs (10μM), 0.5μL MgCl_2_, 0.5μL BSA, forward and reverse primers (see Table 1 for concentrations), and 0.125μL Dreamtaq polymerase. Prior to adding DNA template, DNA samples were briefly vortexed and spun-down to separate the DNA solution from the chelex beads. To each sample, 2μL of DNA template was added. The thermal cycle protocol started at 95°C for 15 min, followed by 35 cycles of denaturation at 94°C for 30 s, annealing at either 51°C or 53°C (see Table 1) for 90 s, extension at 72°C for 90 s, and a final extension at 72°C for 10 minutes. PCR products were stored at 7°C until analysis (within 1 week of extraction). Samples were visually inspected using 2% agarose gel electrophoresis before fragmentation analysis.

**Table 1.**
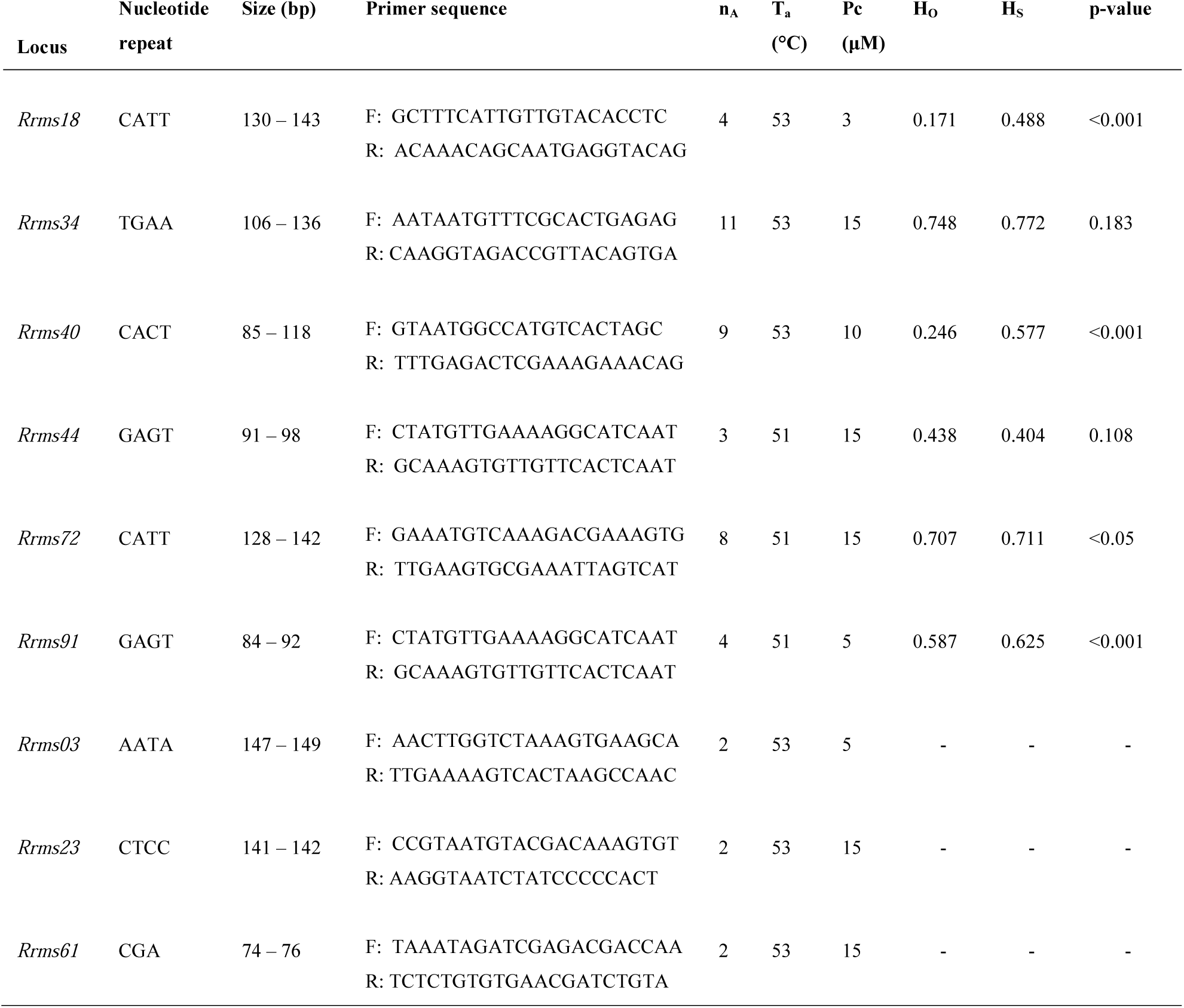
nSSR summary information on each locus. Names, type of repetitive motif, size range of alleles (bp), primer sequences (forward - F, reverse - R), number of alleles (n_A_), annealing temperature (T_a_), primer concentration used in PCR amplification (Pc), and observed (H_O_) and expected (H_S_) heterozygosities, with corresponding p-values.

Primer pairs were labelled with four different fluorescent tags, allowing them to be multiplexed and analysed simultaneously using capillary electrophoresis (ABI PRISM 3100 Genetic Analyzer, Applied Biosystems). Per two amplicons (differently labelled), 1μL of PCR product, 0.3μL of orange dye labelled GeneScan^™^ size standard 500LIZ^™^ (Applied Biosystems), and 10μL formamide was added and denatured before running on the ABI analyzer. Data was visualised, and alleles were scored, in GeneMapper^™^ software (v4.1.1) (Applied Biosystems), after which each automatically scored allele was double-checked by hand. Our nSSRs were defined by a characteristic stutter followed by a peak of at least 450 relative fluorescent units or greater. We further assayed approximately 10% of our samples a second time to check and ensure repeatability of scoring.

### Statistical analyses

With the use of GenoDive v.2.28 [45] that accounts for information gaps by drawing random alleles from the baseline allele frequencies (e.g., missing or null alleles, ensuring no individuals were excluded from analysis), various metrics of genetic diversity were calculated. Beyond calculating the number of alleles per locus (n_A_), we also quantified observed heterozygotes within a subpopulation (i.e., females, fighters, scramblers) (H_O_) and expected frequency of heterozygotes (H_S_) under Hardy-Weinberg equilibrium (HWE) [46], both ranging from 0 to 1. These metrics were then used to calculate the inbreeding coefficient (G_IS_), and determined whether subpopulations departed from HWE (ranging from −1, more heterozygosity than expected, to 1, less heterozygosity than expected). To measure genetic divergence among subpopulations, Wright’s F_ST_ was estimated according to F-statistics defined by Weir & Cockerham [47], whereby the ratio of heterozygosity within the subpopulation is compared to the total population (ranges from 0 - little to no genetic divergence between populations, to 1 - total divergence between subpopulations).

We also performed an analysis of molecular variance (AMOVA) [48] in GenoDive to test for population genetic structure; calculations were performed on four different hierarchical levels (between environments [rich and poor], between subpopulations [sexes and morphs] within environment, among individuals within subpopulations, and within individuals), and gives us insight in the genetic differentiation between these different levels. Statistical significance was evaluated based on 999 random permutations and distances were calculated using the Infinite Alleles Model.

We further subsampled 30 random individuals per group and performed the same analyses with the aim to control for possible artefacts or bias within our analyses stemming from missing data or unequal sampling. Random subsampling and reanalysis was performed 5 times (exemplar represented in Additional File 2, Table S2.1-S2.5).

STRUCTURE analysis (GenoDive v2.28 [45]; STRUCTURE add-in [49]) was additioanlly used to infer genetic clustering using the multilocus nSSR data within populations (rich and poor environment) among the respective subpopulations (i.e. females, scramblers and fighters). This analysis used a Monte Carlo Markov Chain (MCMC) to identify genetically distinct clusters by assigning individuals to K clusters based on assignment probability (Q-value), minimizing departures from HWEand linkage equilibrium. We used a 5 × 10^3^ burn-in, followed by 5 × 10^4^ iterations assuming admixture and correlated allele frequencies without prior population information. We ran 1 to 10 K clusters, with 20 replicates for each cluster. Optimal population clusters were determined according to delta K [50] and bar plot visualisations were compiled using the program STRUCTURE PLOT [51].

## Results

After protocol optimization, we found only 9 of the 16 nSSRs amplified well for our populations, of which 3 loci revealed fixation, and 6 demonstrating both clean/readable peaks and polymorphism across individuals. Thus, these 6 nSSRs were chosen for the genotyping of all remaining individuals.

Across individuals, we had a total of 12.3% missing or null alleles; 3.6% in females, 12.8% in males (25.9% in fighters, 3.7% in scramblers). In the poor environment (19.1%), missing data for females was 5.4%, and for males, 20.5% (34.5% in fighters, and 0.00% in scramblers). In the rich environment (10.2%), missing data for females was 8.65%, and for males, 9.68% (14.1% in fighters, and 6.4% in scramblers). We additionally detected 11 private alleles across 5 loci that differentiated between males and females, and 12 alleles that segregate between the rich (4) or poor (8) environments (Additional File 1, Table S1).

Across all individuals, allelic richness remained low, ranging from 3 to 11 alleles per locus. For four loci, significant deviations from HWE were detected demonstrating an excess of homozygosity present across individuals (Table 1). Deviations from HWE were also detected within our rich and poor environments (Table 2), where rich environments contained significantly lower levels of heterozygosity across all individuals compared to expectation. Poor environments similarly demonstrated lower than expected levels of heterozygosity across all individuals, with the exception of scramblers that were shown to not significantly differ from expectation. These patterns also corresponded to significant levels of inbreeding (G_IS_), with the exception of scramblers in the poor environment.

**Table 2.**
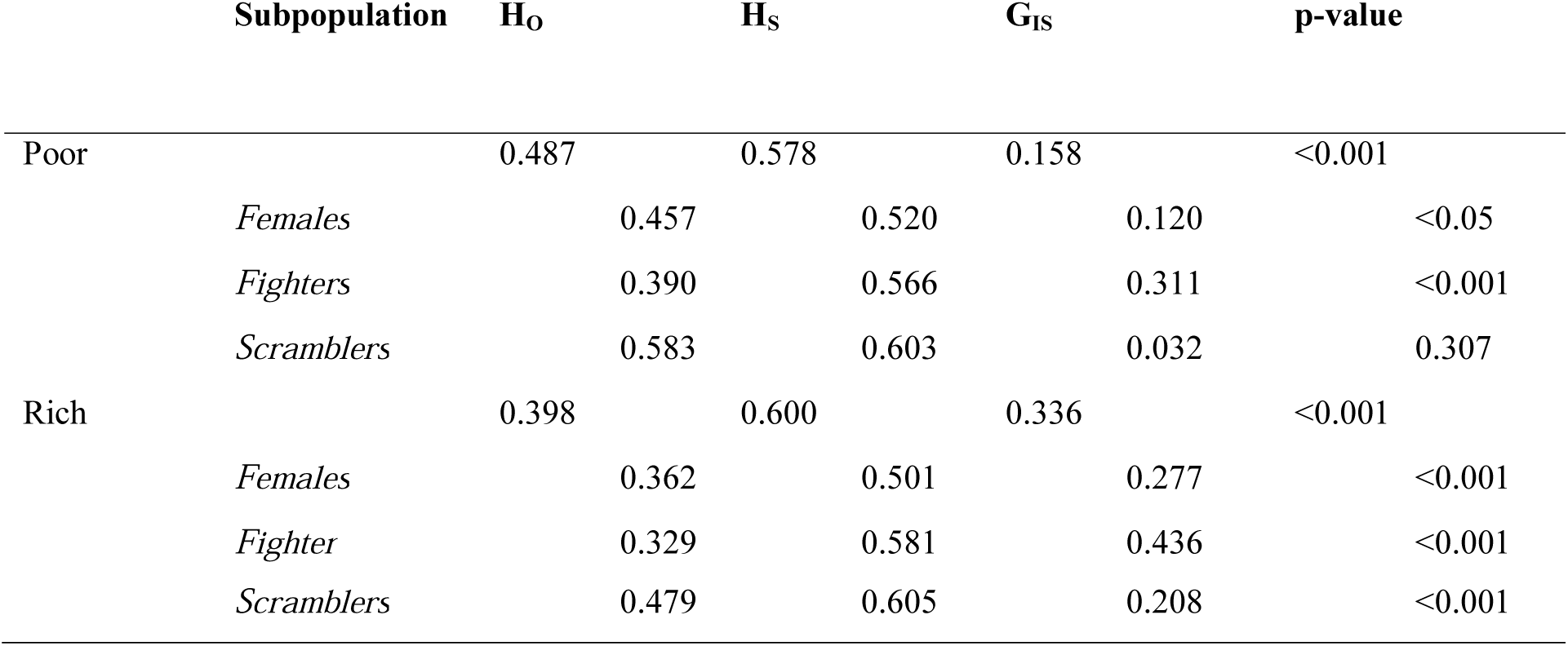
Hardy-Weinberg statistics across environments and subpopulation. Shown are observed (H_O_) and expected (H_S_) heterozygosities, inbreeding coefficient (G_IS_) according to Nei’s statistics (1987), and p-value.

Pairwise genetic differentiation between environments (rich and poor) differed significantly (F_ST_ = 0.109, p<0.001) between the subpopulations (female, fighter and scrambler) within their respective environments (Table 3), with the exception of scramblers compared to females in the poor environment. Although fighters and scramblers significantly differed from each other within both environments (rich and poor), genetic differentiation was approximately three times lower in the poor environment compared to the rich environment (F_ST_ = 0.036, p<0.001, and F_ST_ = 0.102, p<0.001, respectively). These results corroborate the findings that fighters were significantly more genetically divergent compared to scramblers within either environment (Fig. 2).

**Table 3.** Pairwise F_ST_ values for population differentiation. Shown are the genetic differentiation values per subpopulation, Poor (P), and Rich (R) environments, Female (F), Male Fighter (MF), and Male Scrambler (MS) subpopulations. Significant differences are represented by * after Bonferroni correction.

|  | P_F | P_MF | P_MS | R_F | R_MF | R_MS |
| --- | --- | --- | --- | --- | --- | --- |
| **P_F** | -- |  |  |  |  |  |
| **P_MF** | 0.063* | -- |  |  |  |  |
| **P_MS** | 0.024 | 0.036* | -- |  |  |  |
| **R_F** | 0.269* | 0.147* | 0.214* | -- |  |  |
| **R_MF** | 0.085* | 0.103* | 0.109* | 0.213* | -- |  |
| **R_MS** | 0.178* | 0.082* | 0.131* | 0.054* | 0.102* | -- |

**Figure 2.**
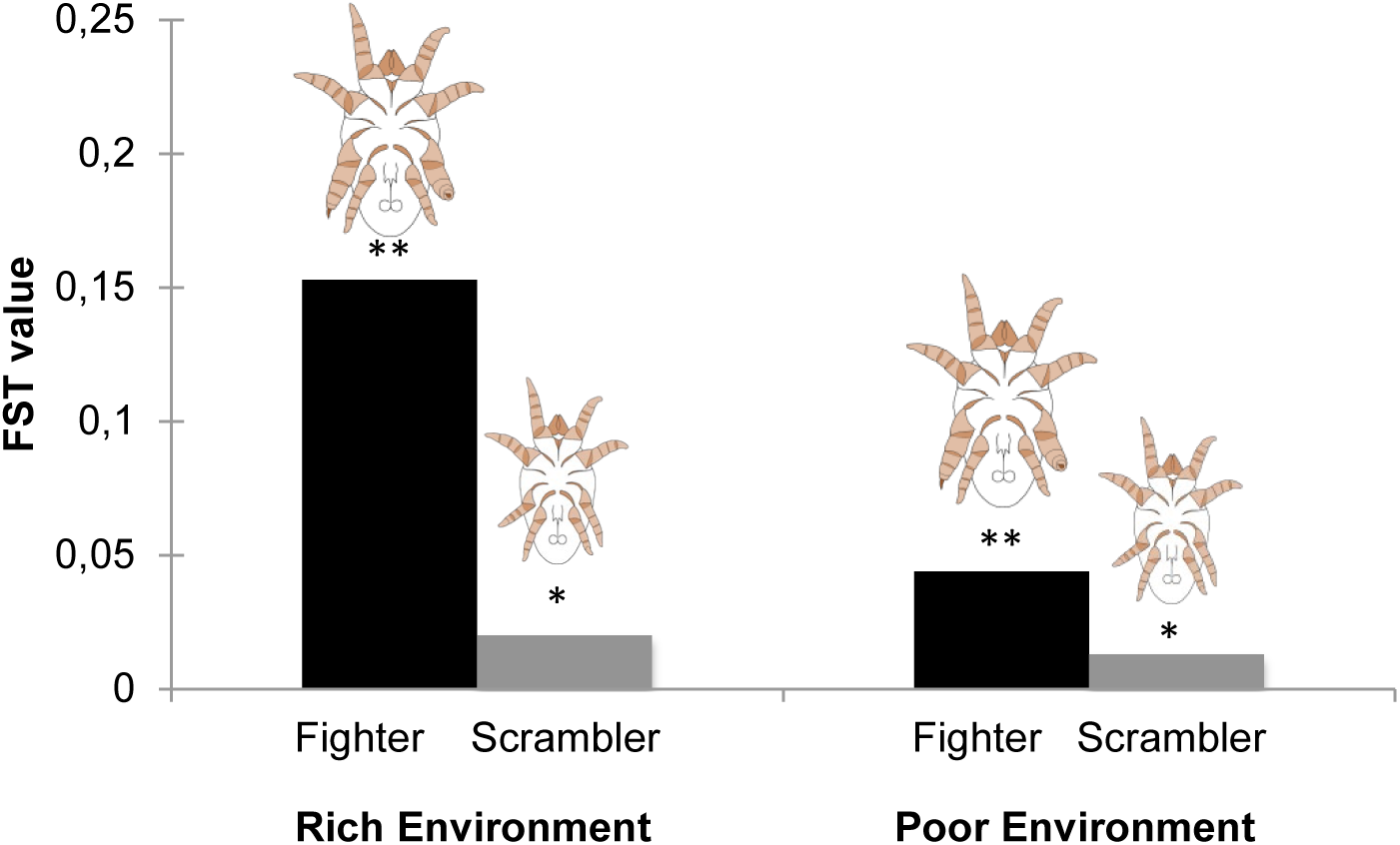
Genetic differentiation (F_ST_) of ART strategies to total population within each environment. Significant differences are represented above bars, * p<0.05, ** p<0.001. ART images kindly supplied by F.T. Rhebergen.

AMOVA analysis (Table 4) showed significant genetic differentiation across subpopulations (females, fighters, and scramblers; F_SC_ = 0.085, p<0.001), between morphs (fighters, and scramblers; F_SC_ = 0.073, p<0.001), and between sexes (females, males; F_SC_ = 0.069, p<0.001). Subpopulations within environment were also genetically different from one another (F_SC_ = 0.085, p<0.001), but the environments (rich and poor) do not differ from the total population (F_CT_ = 0.083, p = 0.206).

**Table 4.** Summary of hierarchical AMOVA. AMOVA including standard deviation (jack-knifing over loci), % of variation, and values of the F-statistic on different levels (between environment, among subpopulation within environment, among individuals within subpopulation, and within individuals), with their corresponding F and p-values. F_CT_ = the proportion of total variance that results from genetic differences among groups, F_SC_ = the proportion of variance among subpopulations within clusters, F_IS_ = the proportion of variance among individuals within subpopulation, F_IT_ = the proportion of variance among individuals within the total population.

| Variance component | SD | Variation (%) | Statistic | F-value | p-value |
| --- | --- | --- | --- | --- | --- |
| *Between environment* | 0.036 | 0.083 | F<sub>CT</sub> | 0.083 | 0.206 |
| *Among subpopulations in environment* | 0.038 | 0.078 | F<sub>SC</sub> | 0.085 | <0.001 |
| *Among individuals in subpopulation* | 0.113 | 0.184 | F<sub>IS</sub> | 0.219 | <0.001 |
| *Within individuals* | 0.120 | 0.655 | F<sub>IT</sub> | 0.345 | <0.001 |

With the exception of locus Rrms 72 demonstrating no significant deviations from HWE (Additional File 2), our subsampled analyses demonstrated near identical results in accordance with our original data set, suggesting any missing/null alleles and unequal sampling within our populations had negligible impact on our results.

Our STRUCTURE analysis demonstrated 2 genetic clusters based on delta K [50] (K=2) best fit our data. Genetic clustering similarly illustrated females and scramblers to disproportionately cluster together compared to fighter individuals that formed their own genetic cluster, although this pattern was more stark in rich compared to poor environments (Fig. 3).

**Figure 3.**
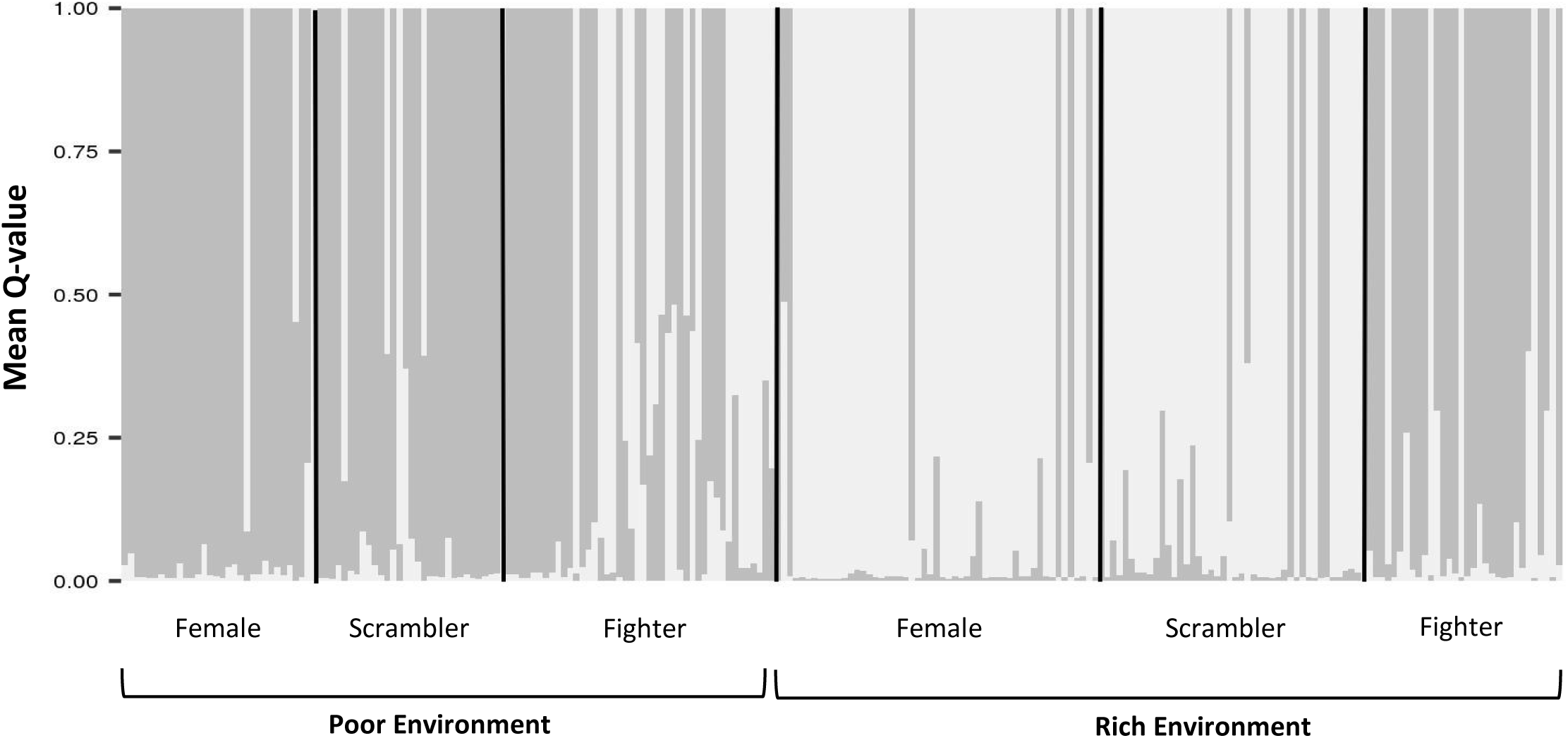
STRUCTURE plot of subpopulation genetic clusters in different environments. STRUCTURE plot illustrating the mean proportional membership (Q-value) of *R. robini* individuals (females, scramblers, fighters) for K=2 across poor and rich environments.

## Discussion

Despite previous formative work focusing on sex- and morph-specific population mean transcriptome patterns in *R. robini* [52, 53], this study is the first of its kind to quantify individual-level genetic diversity in the bulb mite, building a foundation for further genetic quantification investigations for this microscopic organism. Importantly, due to this individual-level approach, the results from this study demonstrate that ARTs in the bulb mite system are associated with genetic diversity, which in-turn is further connected with environmental effects (GEIs). The finding that GEIs underlie the pattern of ARTs is likely to have important repercussions to our understanding of selection in this species, and may help to resolve the previously (but confined) observations for genetic (e.g., [52, 53]) and environmental (e.g., [33, 35, 54]) components operating to mediate male trait expression. GEIs may further help to explain how this polymorphism is maintained within populations over time, notwithstanding often disparate and fluctuating environmental challenges.

### GEIs, genetic context, and the origin of bulb mite ARTs

Counter to our hypothesis for genetic diversity-condition links within our male morphs, we find evidence that large fighters are less genetically variable than their smaller scrambler counterparts. As fighters have been shown to achieve higher reproductive success than scramblers [55], while also being capable of killing conspecifics within populations [56, 57], it is not entirely surprising that these individuals are less genetically diverse simply as a by-product of effective population size reduction [58], and thus genetic erosion. Indeed, both mating monopolization and increased survival likely combine to effectively limit the genetic pool in ensuing fighter offspring. Alternatively, sex-specific effects of inbreeding depression on fitness are also plausible [59], especially in light of high inbreeding consequences on female bulb mite fecundity in general [31]. Previous studies have proposed that the fitness decline of *R. robini* females derived from fighter selection-lines is evidence for intralocus sexual conflict [60]. Our observations that fighters are more inbred than scrambler males, could equally imply that inbred females are less fit and have a higher probability of being purged within populations, similar to life-span and mortality patterns observed in another invertebrate with sex-specific inbreeding depression [61].

A non-mutually exclusive but more adaptive explanation for the origins of the genetic patterns underlying bulb mite ART expression could be their genetic context, or the relation and interaction of genes underpinning this phenotype (epistasis or genetic correlation). Non-additive, epistatic combinations [62, 63] are likely more important than individual genetic components, with pervasive effects from selection to speciation [64]. These genetic interactions have also previously been shown to influence complex traits [65], alter evolutionary trajectories of phenotypes [66], and underlie missing heritability [67].

In the bulb mite specifically, *positive epistasis* could be responsible for fighter expression, such that many alleles in conjunction work in a way that synergistically outperforms their individual contributions to genetically determine the fighter phenotype. Similarly, if many alleles in coordination lead to a less fit phenotype than expected based on their effects in singularity, the process may give rise to a new/alternative phenotype within a population; certain genetic elements combinations may also mask the effects of others (antagonistic epistasis), functionally suppressing the manifestation of high fitness traits (e.g., [68]). The last two aforementioned processes of *negative epistasis* could conceivably produce scramblers within our populations.

Correspondingly, genetic correlations among traits could equally link genetic components together causing similar patterns to the ones we see here. ART-specific genetic correlations have been previously shown in another invertebrate taxa [69], and the breakdown of co-adaptive gene-complexes has been implicated in the adoption of a flexible condition-dependent ART [70], together suggesting that genetic context may be a pervasive, important, but under-investigated facet to ART research. Indeed, markedly distinct genetic patterns among ARTs may be expected owing to the correlational selection for various trait optima combinations between morphs. Ultimately, this correlational selection will result in linkage disequilibrium (opposed and eroded by recombination) having far-reaching evolutionary consequences such as the loss of genetic variation, especially for species frequently undergoing genetic drift through founder effects [7].

Insomuch as complex gene-network for traits are presumed ubiquitous [71], and pleiotropic effects in a single locus for systems necessary to support multi-faceted plasticity (e.g., in morphology, physiology, behaviour) seems dubious [7], it’s likely that heterozygosity in the bulb mite breaks apart genetic elements that require the coordination for the expression of the fighter phenotype, such as specialized developmental trajectories, large body size, aggression, and weaponry. Accordingly, the threshold for fighter development may require a reduction to heterozygosity, such that when heterozygosity within populations decreases, the threshold for fighter expression concomitantly also decreases. Threshold shifts as a response to ART relative fitness would then reflect cryptic genetic variation underlying the translation of the environmental cue to phenotype in a condition-genotype coupling [27]. Future *R. robini* work should aim to assess whether these same GEI patterns are also reflected in natural populations. However, as these broad GEI associations remain consistent between rearing environments, and our rich environment reflects similar natural history responses to that of natural resources (e.g., garlic bulbs [39]), we have no reason to believe that stock and natural populations would differ in their overall patterns of ART genetic context.

### Population-level diversity and the maintenance of ARTs

Considerable variation has been observed in the effects and strength of inbreeding depression, among environments, populations of the same species, and even within sexes (e.g., [61, 72–74]). Our study demonstrates that bulb mites generally lack genetic diversity across individuals, but this pattern could stem from a number of scenarios. For example, in our investigation, near even numbers of scrambler, fighter, and females were collected and compared, yet in reality (stock and wild populations), operational sex-ratios are female skewed ([33, 57], pers. observation), and ART frequencies fluctuate within populations based on environmental milieu [54]. In effect, the average genetic contribution of fighters both within poor and rich environments, compared to the combined contribution of scramblers and females, is likely highly over-represented. Moreover, lab reared populations are known to undergo genetic drift and demonstrate lower than average genetic diversity compared to their wild counterparts [75–77]. However, similar to other species [73], bulb mites may also display a general lack of inbreeding consequences. That said, the combined evidence that fighter phenotypes achieve higher reproductive success than scramblers [55], and that bulb mite ARTs demonstrate some level of heritability [35–37] but no frequency-dependence [34, 55], has continuously raised questions as to how these male polymorphisms are sustained within populations. Certainly the added evidence that fighter phenotypes are also associated with excess homozygosity (this study) further complicates our understanding of how male phenotypic and genetic variation are sustained in this system. Here we link the genetic architecture and life-history parameters of ARTs with oscillating environmental conditions, and suggest that these evolutionary-ecological dynamics may hold the answer.

Previous empirical evidence in bulb mites not only demonstrates that scrambler morphs live longer [78], but importantly, that scrambler-selected lines produce more females that lay larger and more eggs over a longer period of time [79], and are generally more fecund than fighter-selected lines [60]. These morph-specific patterns may help to elucidate why we observed the genetic architecture of scramblers and females to be more similar to each other in contrast to fighters, patterns corroborated in gene expression profiles [52]. Similarly, these reproductive patterns may also help explain how fluctuating environmental conditions, and thus the ensuing shifts in ART frequencies, assist in maintaining genetic diversity within this species. For example, individuals that accumulated deleterious mutations otherwise buffered in optimal conditions (e.g., fighters and possibly female offspring of fighters in the bulb mite) would eventually be purged within poor (presumably stressful) environments (e.g., [74]). This mutation-selection balance could also reduce the genetic differentiation between morphs and sexes, as seen in our bulb mite individuals raised in the poor environment. Certainly, genetic variation in the threshold underlying sensitivity to environmental cues, as assumed in the environmental threshold model [21, 22], would thus cause genetic, and therefore concomitant demographic, oscillation within populations, conceivably facilitating stable population persistence even in the face of spatially or temporally unstable environmental challenges.

Across taxa, processes for the maintenance of genetic diversity are especially significant as they serve as a means for populations to adapt to changing environments and thus play an important role for the survival of a species [80], including reducing its vulnerability to ecological challenges, such as disease or climate change [81]. Whether ARTs buffer populations from excessive inbreeding, and are more likely to evolve in species that routinely encounter boom-bust cycles or environmental perturbations, is certainly a worthy future investigation.

## Conclusion

The complexity, and need for organisms to interact with their environment (to adjust, acclimatize, development, and maximize fitness) implies that genetic context, and thus GEIs, are likely to be pervasive even among plastic phenotypes. Still, the evolution and proximate cause of these phenotypic alternatives are only beginning to be understood. Ultimately, our ability to accurately predict responses to selection based on the genetic variation that maintain thresholds for ARTs, and appreciating the relative genetic and environmental contributions influencing phenotypic expression, is critical to understanding both the breadth and maintenance of within-species variation and a populations capacity to adapt to external adjustments.

## Declarations

### Ethics Approval

In accordance with Dutch law, no ethics approval is required for work conducted on *Rhizoglyphus robini*.

### Consent for Publication

not applicable

### Availability of Data and Material

The datasets generated and/or analysed during the current study are available in the Dryad repository (uploaded upon acceptance).

### Authors’ Contribution

KAS designed the experiment, MRK designed the primers, and RD performed the genetic amplifications and analysis. KAS and RD wrote the majority of the manuscript, KAS, RD and IMS contributed to data interpretation. All authors read, commented on, and approved the final manuscript.

## Acknowledgements

The authors would like to thank Patrick Meirmans for assistance with Genodive, Peter Kuperus for lab support, and Flor Rhebergen for invaluable discussions. We would also like to thank Jan van Arkel and Flor Rhebergen for their bulb mite visualisations. In particular, the authors thanks NWO for funding this research and the anonymous reviewers for their comments on earlier incarnations of the manuscript.

